# Host cellular RNA helicases regulate SARS-CoV-2 infection

**DOI:** 10.1101/2021.06.29.450452

**Authors:** Yasuo Ariumi

## Abstract

Severe acute respiratory syndrome coronavirus 2 (SARS-CoV-2) has largest RNA genome of approximately 30kb among RNA viruses. The DDX DEAD-box RNA helicase is a multifunctional protein involved in all aspects of RNA metabolism. Therefore, host RNA helicases may regulate and maintain such large viral RNA genome. In this study, I investigated the potential role of several host cellular RNA helicases in SARS-CoV-2 infection. Notably, DDX21 knockdown markedly accumulated intracellular viral RNA and viral production, as well as viral infectivity of SARS-CoV-2, indicating that DDX21 strongly restricts the SARS-CoV-2 infection. As well, MOV10 RNA helicase also suppressed the SARS-CoV-2 infection. In contrast, DDX1, DDX5, and DDX6 RNA helicases were required for SARS-CoV-2 replication. Indeed, SARS-CoV-2 infection dispersed the P-body formation of DDX6 and MOV10 RNA helicases as well as XRN1 exonuclease, while the viral infection did not induce stress granule formation. Accordingly, the SARS-CoV-2 nucleocapsid (N) protein interacted with DDX6, DDX21, and MOV10 and disrupted the P-body formation, suggesting that SARS-CoV-2 N hijacks DDX6 to utilize own viral replication and overcomes their anti-viral effect of DDX21 and MOV10 through as interaction with host cellular RNA helicase. Altogether, host cellular RNA helicases seem to regulate the SARS-CoV-2 infection.

**Importance:** SARS-CoV-2 has large RNA genome of approximately 30kb. To regulate and maintain such large viral RNA genome, host RNA helicases may involve in SARS-CoV-2 replication. In this study, I have demonstrated that DDX21 and MOV10 RNA helicases limit viral infection and replication. In contrast, DDX1, DDX5 and DDX6 are required for the SARS-CoV-2 infection. Interestingly, the SARS-CoV-2 infection disrupted P-body formation and attenuated or suppressed stress granule formation. Thus, SARS-CoV-2 seems to hijack host cellular RNA helicases to play a proviral role by facilitating viral infection and replication and, by suppressing host innate immune system.

## Introduction

The DDX DEAD-box RNA helicase family, which is an ATPase-dependent RNA helicase, is a multifunctional protein involved in all aspects of RNA life cycle, including transcription, mRNA splicing, RNA transport, translation, ribosome biogenesis, RNA decay, and viral infection (1–6). DDX3 RNA helicase has two homologs termed DDX3X and DDX3Y, which were located on X and Y chromosome. Actually, DDX3 involves in translation, transcription, cell cycle, tumorigenesis (oncogenic and tumor suppressor function), viral infection and innate immunity (7). Indeed, DDX3 is known to be a component of anti-viral innate immune signaling pathway and contributes to induce anti-viral mediators, such as type I interferon and interferon regulatory factor 3 (IRF3). On the other hand, DDX3 is required for both human immunodeficiency virus type 1 (HIV-1) and hepatitis C virus (HCV) infection and replication (7–12). In addition, we identified DDX1, DDX5, DDX17, and DDX21 RNA helicases as HIV-1 Rev-interacting proteins that enhanced the Rev-dependent RNA export function (8). autoimmune antibody from a patient with watermelon stomach disease (13). DDX21 interacts with c-Jun and acts as a co-factor for c-Jun-mediated transcription (14). DDX21 regulates transcription and ribosomal RNA processing (15). DDX6 (Rck/p54) RNA helicase predominantly localizes in the discrete cytoplasmic foci called the processing (P)-body. DDX6 interacts with an initiation factor eIF-4E to repress the translational activity of mRNA (16). DDX6 regulates the activity of the decapping enzymes DCP1 and DCP2 and directly interacts with Argonaute-1 (Ago1) and Ago2 in microRNA (miRNA)-induced silencing complex (miRISC) and involves in RNA-mediated gene silencing (RNAi). Thus, P-body is an aggregate of translationally repressed mRNA associated with the translation repression and mRNA decay machinery (17, 18). In fact, DDX6 negatively regulate HIV-1 gene expression by preventing HIV-1 mRNA association with polysomes (19). In contrast, DDX6 is required for HCV replication and HIV-1 capsid assembly (12, 20–22). Furthermore, the SF1 RNA helicase Moloney leukemia virus 10 (MOV10) also localizes in P-body. MOV10 is incorporated into HIV-1 virion and acts as an antiviral factor (23–25).

In addition, host cells contain another type of RNA granule called stress granule (SG) (26, 27). SGs are aggregates of untranslating mRNAs in conjunction with translation initiation factors (eIF4E, eIF3, eIF4A, eIFG, and PABP), the 40S ribosomal subunits, and several RNA-binding proteins including poly(A)-binding protein (PABP), GTPase-activating protein (SH3 domain) binding protein 1 (G3BP1), Ataxin-2 (ATX2), T-cell intracellular antigen-1 (TIA-1), and TIA-R, and SGs regulate mRNA translation and decay, as well as proteins involved in various aspects of mRNA metabolisms. SGs are cytoplasmic phase-dense structures that occur in cells exposed to various environmental stress, including heat, arsenite, viral infection, oxidative condition, UV irradiation, and hypoxia. SGs and P-bodies physically interact and mRNPs may shuttle between two compartments (17, 18, 26, 27). On the other hand, several viruses targets SGs and P-bodies for the viral replication (12, 17). We previously demonstrated that HCV infection disrupted P-body formation of DDX6, Lsm1, Xrn1, and Ago2 and induced SGs formation of G3BP1, PABP, and ATX2 (12). HCV hijacks the P-body and stress granule components around lipid droplets (LDs) to utilize own viral replication (7, 12).

Severe acute respiratory syndrome coronavirus 2 (SARS-CoV-2), which is the causative agent of coronavirus disease 2019 (COVID-19), infects more than 100 million people worldwide and remains a global public health problem (28–32). SARS-CoV-2 is an enveloped, positive-stranded RNA virus with largest RNA genome of ∼30kb among RNA viruses and belongs to the *Coronaviridae* family, including SARS-CoV and Middle East respiratory syndrome coronavirus (MERS-CoV). SARS-CoV-2 encodes 16 non-structural proteins (Nsp1-Nsp16), nine putative accessory proteins, and four structural proteins, spike (S), envelope (E), membrane (M), and nucleocapsid (N) proteins (33). The N protein forms a helical ribonucleocapsid complex with viral genomic RNA and interacts with viral membrane protein M during assembly of virion. The S protein allows the virus to attach to and fuse with host cell membrane for viral entry. Angiotensin converting enzyme 2 (ACE2) acts as the receptor for SARS-CoV-2 (34–36). To regulate and maintain such large viral RNA genome, host RNA helicases may involve in SARS-CoV-2 replication. To address this issue, in this study, I investigated the potential effect of host cellular RNA helicases on SARS-CoV-2 infection.

## Materials and Methods

### Cell culture

293T, HepG2, HEK293T ACE2 (SL221, GeneCopoeia Inc., Rockville, MD, USA), and VeroE6 TMPRSS2 cells (a gift from Drs. Shutoku Matsuyama and Makoto Takeda, National Institute of Infectious Diseases, Tokyo, Japan) (37) were cultured in Dulbecco’s modified Eagle’s medium (DMEM; Life Technology, Carlsbad, CA, USA) supplemented with 10% fetal bovine serum (FBS). CACO-2 human colon carcinoma cells (RCB0988; RIKEN BioResource Research Center, Tsukuba, Ibaraki, Japan) were cultured in DMEM supplemented with 20% FBS.

### Plasmid construction

To construct pcDNA3-HA-MOV10, a DNA fragment encoding MOV10 was amplified from HuH-7 cDNA by PCR using KOD-Plus DNA polymerase (TOYOBO, Osaka, Japan) and the following pairs of primers: 5’-CGGGATCCAAGATGCCCAGTAAGTTCAGCTG-3’ (Forward), 5’-CCGCTCGAGTCAGAGCTCATTCCTCCACTCTG-3’ (Reverse). The obtained DNA fragments were subcloned into either the *Bam*HI-*Not*I sites of the pcDNA3-HA vector, and the nucleotide sequences were determined by Sanger sequencing. We previously described pHA-DDX3 (8–12), pcDNA3-HA-DDX1 and pcDNA3-HA-DDX6 (9). pcDNA3.1 SARS-CoV-2 N was a gift from Dr. Jeremy Luban (Addgene plasmid #158079; http://n2t.net/addgene:158079; RRID:Addgene_158079) (38).

### Western blot analysis

Cells were lysed in buffer containing 50 mM Tris-HCl (pH 8.0), 150 mM NaCl, 4 mM EDTA, 1% Nonidet (N) P-40, 0.1% sodium dodecyl sulfate (SDS), 1 mM dithiothreitol (DTT) and 1 mM phenylmethylsulfonyl fluoride (PMSF). Supernatants from these lysates were subjected to SDS-polyacrylamide gel electrophoresis, followed by immunoblot analysis using anti-SARS-CoV-2 Spike (GTX632604 [1A9]; GeneTex, Irvine, CA, USA), anti-DDX3 (A300-474A; Bethyl Lab, Inc., Montgomery, TX, USA), anti-DDX1 (A300-521A; Bethyl Lab), anti-DDX5 (A300-523A; Bethyl Lab), anti-DDX6 (A300-460A; Bethyl Lab), anti-DDX21 (A300-627A; Bethyl Lab), anti-MOV10 (A301-571A; Bethyl Lab), anti-G3BP1 (A302-033A; Bethyl Lab), or anti-β-actin (A5441; Sigma, Saint Louis, MI, USA). We used peroxidase-conjugated Donkey anti-Rabbit IgG (H+L) (Jackson ImmunoResearch, West Grove, PA, USA) or Amersham ECL peroxidase-linked Sheep anti-Mouse IgG (GE Healthcare Bio-Sciences, Uppsala, Sweden) as secondary antibodies. The proteins were detected by using Western Lightning Plus ECL, enhanced chemiluminescence substrate (PerkinElmer, Waltman, MA) and ImageQuant LAS4000 system (GE healthcare Bio-Sciences).

### RNA interference

Oligonucleotides with the following sense and antisense sequences were used for the cloning of short hairpin RNA (shRNA)-encoding sequences targeted to DDX1, DDX21, or MOV10 in a lentiviral vector: DDX1, 5’GATCCCCGGAGATGTAAGATTCTTGATTCAAGAGATCAAGAATCTTACATCTCCTTTTTGGAAA3’ (sense), 5’-AGCTTTTCCAAAAAGGAGATGTAAGATTCTTGATCTCTTGAATCAAGAATCTTACATCTCCGGG-3’ (antisense), DDX21, 5’GATCCCCGGAGCATCTGGCTATTAAGTTCAAGAGACTTAATAGCCAGATGCTCCTTTTTGGAAA3’ (sense), 5’-AGCTTTTCCAAAAAGGAGCATCTGGCTATTAAGTCTCTTGAACTTAATAGCCAGATGCTCCGGG-3’ (antisense), MOV10, 5’GATCCCCGCTGACCTTCAAGGTGAACTTCAAGAGAGTTCACCTTGAAGGTCAGCTTTTTGGAAA3’ (sense), 5’-AGCTTTTCCAAAAAGCTGACCTTCAAGGTGAACTCTCTTGAAGTTCACCTTGAAGGTCAGCGGG-3’ (antisense). The oligonucleotides above were annealed and subcloned into the *Bgl*II-*Hin*dIII site, downstream from an RNA polymerase III promoter of pSUPER (39), to generate pSUPER-DDX1, pSUPER-DDX21, and pSUPER-MOV10, respectively. To construct pLV-shDDX1, pLV-shDDX21, or pLV-shMOV10, the *Bam*HI-*Sal*I fragments of the corresponding pSUPER plasmids were subcloned into the *Bam*HI-*Sal*I site of pRDI292, an HIV-1-derived self-inactivating lentiviral vector containing a puromycin resistance marker allowing for the selection of transduced cells (40). We previously described pLV-shDDX3, pLV-shDDX5, and pLV-shDDX6 (11, 12, 41).

### Lentiviral vector production

The vesicular stomatitis virus (VSV)-G-pseudotyped HIV-1-based vector system has been described previously (42, 43). The lentiviral vector particles were produced by transient transfection of the second-generation packaging construct pCMVΔR8.74 (42, 43) and the VSV-G-envelope-expressing plasmid pMD2G as well as pRDI292 into 293T cells with TransIT-LT1 transfection reagent (Mirus Bio LLC, Madison, WI, USA).

### Immunoprecipitation

Cells were lysed in buffer containing 50 mM Tris-HCl (pH 8.0), 150 mM NaCl, 4 mM EDTA, 0.1% NP-40, 10 mM NaF, protease inhibitor cocktail (Nacalai Tesque, Kyoto, Japan). Lysates were pre-cleared with 30 μl of protein-G-Sepharose (GE Healthcare Bio-Sciences). Pre-cleared supernatants were incubated with 5 μg of anti-SARS-CoV-2 nucleocapsid antibody mixture (GTX135357 and GTX632269 [6H3]; GeneTex), anti-DDX21 (A300-627A; Bethyl Lab), anti-DDX6 (A300-460A; Bethyl Lab), or anti-MOV10 (A301-571A; Bethyl Lab) antibody at 4°C for 1 hour (h). Following absorption of the precipitates on 30 μl of protein-G-Sepharose resin for 1 h, the resin was washed four times with 700 μl of lysis buffer. Proteins were eluted by boiling the resin for 5 min in 2× Laemmli sample buffer. The proteins were then subjected to SDS-PAGE, followed by immunoblot analysis using anti-SARS-CoV-2 nucleocapsid (GTX632269 [6H3]), anti-DDX21, anti-DDX6 or anti-MOV10 antibody.

### Immunofluorescence and confocal microscopic analysis

Cells were grown on Lab-Tek 2 well chamber slide (Nunc, Thermo) at 2×10^4^ cells per well. The cells were fixed in 3.6% formaldehyde in phosphate-buffered saline (PBS), permeabilized in 0.1% NP-40 in PBS at room temperature, and incubated with anti-HA antibody (3F10; F. Hoffmann-La Roche AG, Basel, Switzerland) at a 1:300 dilution in PBS containing 3% bovine serum albumin (BSA) at 37°C for 30 min. We also used anti-DDX6 (A300-460A; Bethyl Lab), anti-XRN1 (A300-443A; Bethyl Lab), anti-G3BP1 (A302-033A; Bethyl) anti-DDX21 (A300-627A; Bethyl Lab), anti-MOV10 (A301-571A; Bethyl Lab), anti-DDX5 (A300-523A; Bethyl Lab), or anti-SARS-CoV-2 nucleocapsid (ab273434 [6H3]; Abcam or GTX632269 [6H3]; GeneTex) antibodies as primary antibodies. Cells were then stained with Donkey anti-mouse IgG (H+L) secondary antibody, Alexa Fluor 594 conjugate and/or Donkey anti-rabbit or anti-rat IgG (H+L) secondary antibody, Alexa Fluor 488 conjugate (Thermo Fisher Scientific Inc., Waltham, MA, USA) at a 1:300 dilution in PBS containing 3% BSA at 37°C for 30 min. Nuclei were stained with DAPI (4’, 6’-diamidino-2-phenylindole). Following washing 3 times in PBS, the coverslides were mounted on slides using SlowFade Gold antifade reagent (Life Technology). Samples were analyzed under a confocal laser-scanning microscope (FV1200; Olympus, Tokyo, Japan).

### Virus infection

In this study, I used SARS-CoV-2 2019-nCoV/Japan/TY/WK-521/2020 (GenBank accession no. LC522975.1) isolated from a patient in Japan. The virus was amplified in VeroE6 TMPRSS2 cells in high glucose DMEM supplemented 10%FBS, incubated at 37°C in 5% CO_2_ during 2 days of infection. Virus titers were performed by the tissue culture infectious dose at 50% (TCID_50_/ml) and the virus stocks (2.7×10^8^ TCID_50_/ml) kept in −80°C deep freezer until use. All virus infection studies were performed in biosafety level 3 (BSL3) facility. For infection experiments with SARS-CoV-2 virus, HEK293T ACE2 cells (2×10^5^ cells/well) were plated onto 6-well plates and cultured for 24 h. Then, cells were infected at a multiplicity of infection (MOI) of 0.5. Actually, 1μl of culture supernatants of SARS-CoV-2-infected VeroE6 TMPRSS2 cells (2.7×10^8^ TCID_50_/ml) were inoculated. The culture supernatants were collected at 72 h post-infection and the levels of the nucleocapsid (N) protein were determined by enzyme-linked immunosorbent assay using RayBio COVID-19/SARS-CoV-2 Nucleocapsid protein ELISA kit (RayBiotech, Norcross, GA, USA) and xMark microplate spectrometer (Bio-Rad). Total RNA was also isolated from the infected cellular lysates using RNeasy mini kit (Qiagen, Hilden, Germany) for analysis of intracellular SARS-CoV-2 RNA. The extracellular SARS-CoV-2 RNA in the culture supernatant was analyzed using SARS-CoV-2 direct detection RT-qPCR kit (RC300A; TaKaRa-Bio, Shiga, Japan). The infectivity of SARS-CoV-2 in the culture supernatants was determined at 48 h post-infection. The SARS-CoV-2-infected cells were detected using anti-SARS-CoV-2 Spike (GTX632604 [1A9]; GeneTex, Irvine, CA, USA) or anti-SARS-CoV-2 nucleocapsid (ab273434 [6H3]; Abcam or GTX632269 [6H3]; GeneTex) antibodies.

### Real-time RT-PCR

Total RNA was isolated using RNeasy Mini kit (Qiagen), and cDNAs were synthesized using SARS-CoV-2 nucleocapsid (N) reverse primer N660R (5’-AGCAAGAGCAGCATCACCGCCATTGCCAGC-3’) and M-MLV reverse transcriptase (Invitrogen). Then, real-time RT-PCR was performed with *SARS-CoV-2 N* primer sets and SYBR Premix Ex Taq II (TaKaRa-Bio) using a LightCycler Nano (Roche) with 40 cycles. Primer sequences are as follows: 5’-ATGCTGCAATCGTGCTACAA-3’ (Forward), 5’-GACTGCCGCCTCTGCTC-3’ (Reverse) for *SARS-CoV-2 N*.

### Statistical analysis

A statistical comparison of the RNA levels of SARS-CoV-2 between the knockdown cells and the control cells was performed by using the Student t test. Two-sided P values of less than 0.05 were considered statistically significant. Data are shown as mean ± standard error of the mean (SEM) three independent experiments.

## Results

### DDX21 RNA helicase restricts SARS-CoV-2 infection

To study the potential role of host cellular RNA helicases in SARS-CoV-2 infection, I first used lentiviral vector-mediated RNA interference to stably knockdown DDX1, DDX3, DDX5, DDX21 or MOV10 RNA helicase in HEK293T ACE2 cells, which constitutively express ACE2, a SARS-CoV-2 receptor (34–36). Then, I used puromycin-resistant pooled cells 10 days after the lentiviral transduction in all experiments. Western blot showed a very effective knockdown of each RNA helicase in HEK293T ACE2 cells transduced with lentiviral vectors expressing the corresponding shRNAs (Figure 1A). Importantly, shRNAs did not affect cell viabilities (data not shown). We next examined the level of intracellular SARS-CoV-2 RNA in these knockdown cells 72 h after SARS-CoV-2 infection. Real-time LightCycler RT-PCR analysis demonstrated that the accumulation of SARS-CoV-2 RNA was significantly suppressed in DDX1, or DDX5 knockdown cells (Figure 1B), indicating that both DDX1 and DDX5 are required for SARS-CoV-2 lifecycle. In contrast, I noticed that the level of intracellular RNA was strongly enhanced in DDX21 knockdown cells (approximately 4,000-fold) compared with that of the control cells (Figure 1B), suggesting that DDX21 strongly restricts SARS-CoV-2 infection. In addition, MOV10 RNA helicases also suppressed the SARS-CoV-2 infection. Consistent with these findings, Western blot analysis also showed that intracellular SARS-CoV-2 spike protein expression was markedly enhanced in the DDX21 knockdown cells (Figure 1C). To further confirm the anti-viral effect of DDX21, I examined the DDX21 knockdown CACO-2 human colon cancer cells and HepG2 human hepatoma cells (Figure 1D). CACO-2 cells express endogenous ACE2 and are susceptible to SARS-CoV-2 infection. The DDX21 knockdown demonstrated enhanced intracellular SARS-CoV-2 RNA in both CACO-2 and HepG2 cells (Figure 1D). Thus, DDX21 seems to restrict

**Figure 1.**
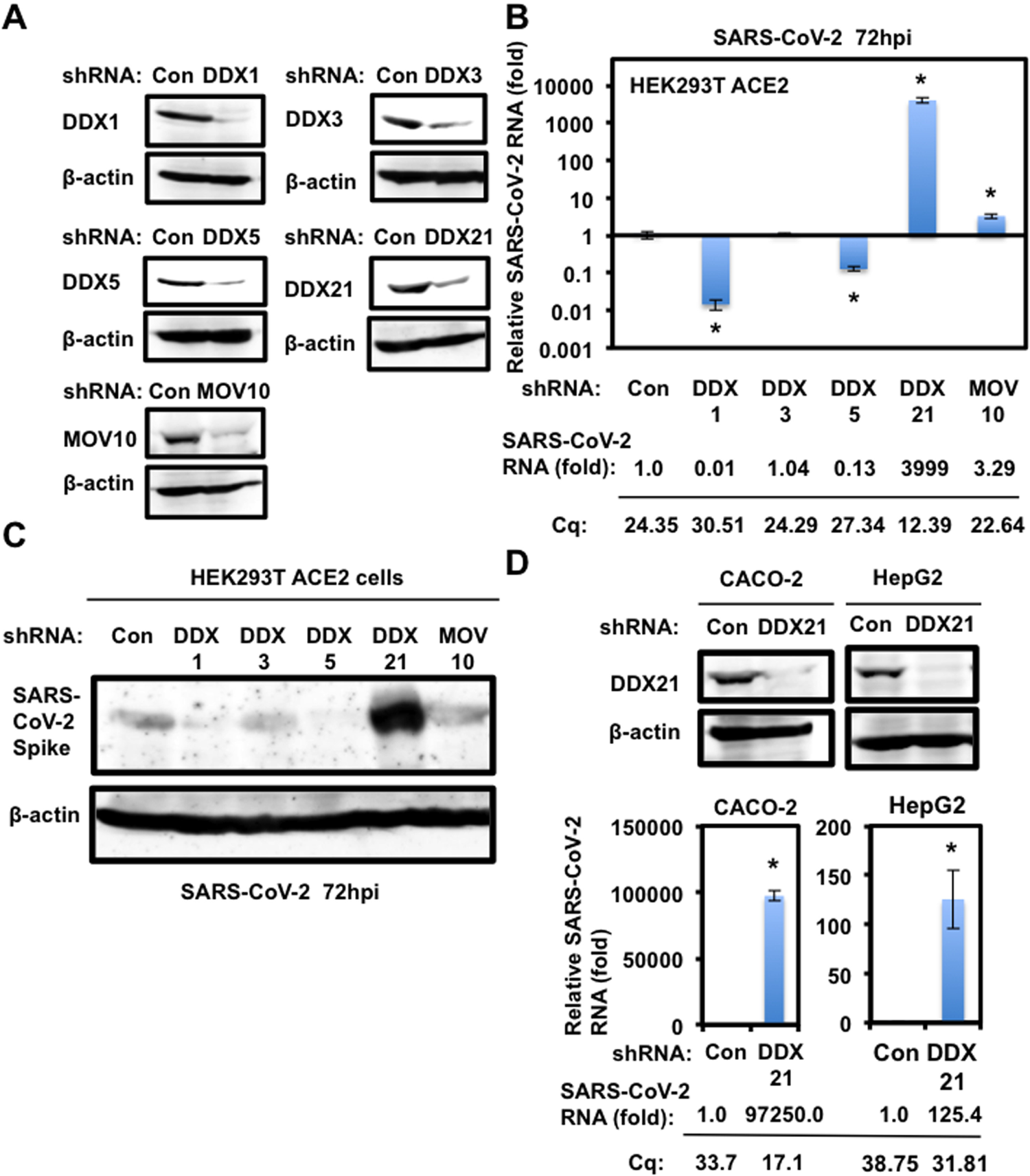
DDX21 restricts SARS-CoV-2 infection. (**A**) Inhibition of host cellular RNA helicase protein expression by the shRNA-producing lentiviral vector. The results of Western blot analysis of cellular lysates with anti-DDX1 (A300-521A), anti-DDX3 (A300-474A), anti-DDX6 (A300-460A), anti-DDX21 (A300-627A), anti-MOV10 (A301-571A), or anti-β-actin antibody are shown. (**B**) The level of intracellular SARS-CoV-2 RNA in the cells at 72 h post-infection at an MOI of 0.5 was monitored by real-time LightCycler RT-PCR. Results from three independent experiments are shown. The level of SARS-CoV-2 RNA in each knockdown cells was calculated relative to the level in HEK293T ACE2 cells transduced with a control lentiviral vector (Con). Asterisk indicates significant differences compared to the control cells. *P<0.05. Cq, quantitation cycle. (**C**) SARS-CoV-2 spike protein expression levels in each knockdown cells. The results of the Western blot analysis of cellular lysates with anti-SARS-CoV-2 Spike (GTX632604 [1A9]) or anti-β-actin antibody in the SARS-CoV-2-infected HEK293T ACE2 cells at 72 h post-infection at an MOI of 0.5 are shown. (**D**) DDX21 restricts SARS-CoV-2 infection in CACO-2 and HepG2 cells. Inhibition of endogenous DDX21 protein expression by the shRNA-producing lentiviral vector. The results of Western blot analysis of cellular lysates with anti-DDX21 or anti-β-actin antibody in CACO2 or HepG2 cells are shown. The level of intracellular SARS-CoV-2 RNA in the cells at 72 h post-infection at an MOI of 0.5 was monitored by real-time LightCycler RT-PCR. Results from three independent experiments are shown. The level of SARS-CoV-2 RNA in DDX21 knockdown cells was calculated relative to the level in HEK293T ACE2 cells transduced with a control lentiviral vector (Con). Asterisk indicates significant differences compared to the control cells. *P<0.05.

### SARS-CoV-2 infection

We next examined the levels of extracellular SARS-CoV-2 nucleocapsid (N) protein and the level of extracellular SARS-CoV-2 RNA as well as the infectivity of SARS-CoV-2 in the culture supernatants in these knockdown cells 72 h after SARS-CoV-2 infection. The results showed that the release of SARS-CoV-2 N and the accumulation of extracellular SARS-CoV-2 RNA were significantly enhanced in DDX21 knockdown cells (Figure 2A and 2B). In this context, and the virus titer and the infectivity of SARS-CoV-2 in the culture supernatants were also significantly enhanced in these knockdown cells (Figure 2C and 2D). Importantly, I noticed that the viral titer in the culture supernatants of DDX21 knockdown cells was elevated approximately 10,000-fold compared with that of control cells, while the level of extracellular SARS-CoV-2 N protein and extracellular SARS-CoV-2 RNA in the supernatants of DDX21 knockdown cells were elevated only 5-fold compared with that of control cells (Figure 2A-2D), suggesting that DDX21 suppresses the viral RNA replication, the viral production as well as the viral infectivity. Then, I examined subcellular localization of SARS-CoV-2 N protein in the control and DDX21 knockdown HEK293T ACE2 cells 48 h after inoculation of SARS-CoV-2. The SARS-CoV-2 N protein predominantly localized in cytoplasm however, there was no clear difference between the control and DDX21 knockdown cells (Figure 2E). Furthermore, I examined the subcellular localization of endogenous DDX21 and SARS-CoV-2 N in both VeroE6 TMPRSS2 and HEK293T ACE2 cells 24 h after inoculation of SARS-CoV-2. DDX21 predominantly localized in nucleoli and SARS-CoV-2 N localized in cytoplasm in SARS-CoV-2-infected cells (Figure 2F).

**Figure 2.**
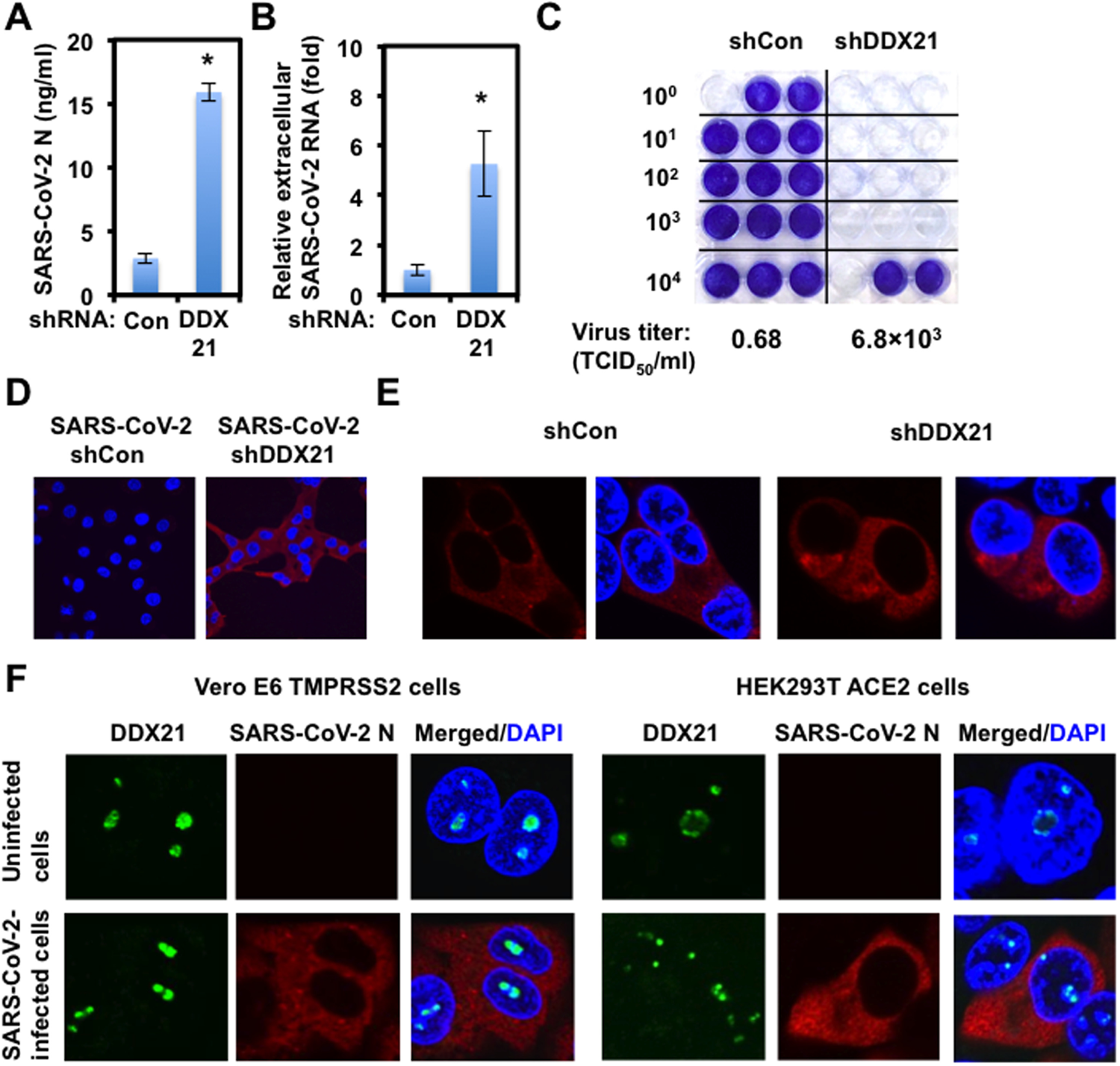
Characterization of anti-viral effect of DDX21. (**A**) DDX21 suppresses SARS-CoV-2 production. The levels of extracellular SARS-CoV-2 N protein in the culture supernatants from the DDX21 knockdown HEK293T ACE2 cells 72 h after inoculation of SARS-CoV-2 at an MOI of 0.5 were determined by ELISA. Experiments were done in triplicate and columns represent the mean SARS-CoV-2 N protein levels. Asterisk indicates significant differences compared to the control cells. *P<0.05. (**B**) DDX21 inhibits the level of extracellular SARS-CoV-2 RNA. The level of extracellular SARS-CoV-2 RNA in the culture supernatants from the DDX21 knockdown HEK293T ACE2 cells 72 h after inoculation of SARS-CoV-2 at an MOI of 0.5 were monitored by real-time LightCycler RT-PCR. Results from three independent experiments are shown. The level of SARS-CoV-2 RNA in DDX21 knockdown cells was calculated relative to the level in HEK293T ACE2 cells transduced with a control lentiviral vector (Con). Asterisk indicates significant differences compared to the control cells. *P<0.05. (**C**) The virus titer of SARS-CoV-2 in the culture supernatants from the DDX21 knockdown HEK293T ACE2 cells 72 h after inoculation of SARS-CoV-2. Naïve VeroE6 TMPRSS2 cells were seeded in 24-well plates at 5×10^4^ cells per well and then infected the next day with the indicated serial 10-fold dilutions of culture supernatants. The cells were stained with 0.6% Coomassie brilliant blue in 50% methanol and 10% acetate at 72 h post-infection were monitored for cytopathic effect (CPE). The virus titer was determined by the tissue culture infectious dose at 50% (TCID_50_/ml). (**D**) The infectivity of SARS-CoV-2 in the culture supernatants from the control or DDX21 knockdown HEK293T ACE2 cells 72 h after inoculation of HCV-JFH1 was compared by immunofluorescence. Naïve VeroE6 TMPRSS2 cells were plated on Lab-Tek 2 well chamber slide at 2×10^4^ cells per well. Next day, 1μl of culture supernatants of SARS-CoV-2-infected control or DDX21 knockdown HEK293T ACE2 cells were inoculated. The cells were fixed at 24 h post-infection and stained with anti-SARS-CoV-2 nucleocapsid (ab273434 [6H3]). Cells were then stained with Donkey anti-mouse IgG (H+L) secondary antibody, Alexa Fluor 594 conjugate. Images were visualized using confocal laser scanning microscopy. Nuclei were stained with DAPI (blue). (**E**) Subcellular localization of SARS-CoV-2 N protein in control or DDX21 knockdown HEK293T ACE2 cells 24 h after inoculation of SARS-CoV-2. The cells were stained with anti-SARS-CoV-2 nucleocapsid. (**F**) Subcellular localization of endogenous DDX21 and SARS-CoV-2 N protein in VeroE6 TMPRSS2 or HEK293T ACE2 cells 24 h after inoculation of SARS-CoV-2. The cells were stained with anti-SARS-CoV-2 nucleocapsid and anti-DDX21 (A300-627A) antibodies. Cells were then stained with Donkey anti-rabbit IgG (H+L) secondary antibody, Alexa Fluor 488 conjugate and Donkey anti-mouse IgG (H+L) secondary antibody, Alexa Fluor 594 conjugate. The two-color overlay images are also exhibited (Merged).

### SARS-CoV-2 disrupts P-body formation and hijacks host cellular RNA helicases

To investigate the potential role of P-body components in SARS-CoV-2 infection, I first examined the alteration of subcellular localization of DDX6 RNA helicase by SARS-CoV-2 infection using confocal laser scanning microscopy. As we previously described (12), DDX6, predominantly localized in the evident cytoplasmic foci termed P-bodies in both uninfected naïve VeroE6 TMPRSS2 and HEK293T ACE2 cells (Figure 3). Notably, SARS-CoV-2 infection disrupted the P-body formation of DDX6 in both cells at 24 h post-infection (Figure 3). Indeed, endogenous DDX6 dispersed in the cytoplasm and colocalized with SARS-CoV-2 N in response to the SARS-CoV-2 infection (Figure 3 and 4A). Therefore, we further examined whether or not SARS-CoV-2 disrupts the P-body formation of other P-body components, including MOV10 RNA helicase and the 5’-3’ exonuclease Xrn1. As expected, SARS-CoV-2 infection disrupted the P-body formation of MOV10 and XRN1 (Figure 3). I also observed that most of DDX6 still formed intact P-bodies at earlier time (6 h post-infection) (Figure 4A). Then, I noticed that P-body formation of DDX6 began to be disrupted at 24 h post-infection (Figure 4A).

**Figure 3.**
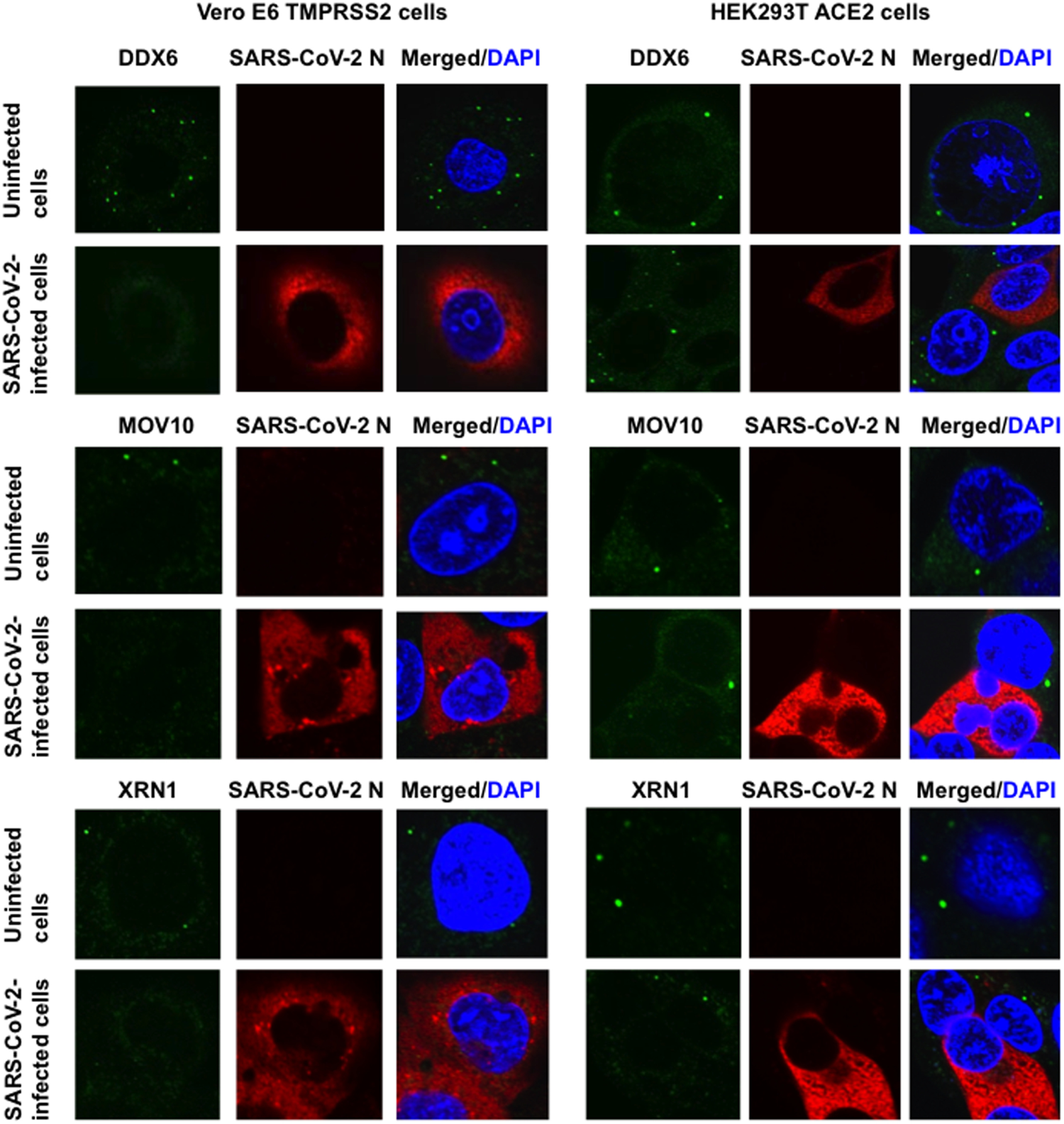
SARS-CoV-2 disrupts the P-body formation. Uninfected VeroE6 TMPRSS2 or HEK293T ACE2 cells and their SARS-CoV-2-infected cells at 24 h post-infection were stained with anti-SARS-CoV-2 nucleocapsid (ab273434 [6H3]) and anti-DDX6 (A300-460A) antibodies. The cells were also stained with anti-SARS-CoV-2 nucleocapsid and either anti-Xrn1 (A300-443A) or anti-MOV10 (A301-571A) antibodies. Cells were then stained with Donkey anti-rabbit IgG (H+L) secondary antibody, Alexa Fluor 488 conjugate and Donkey anti-mouse IgG (H+L) secondary antibody, Alexa Fluor 594 conjugate. Images were visualized using confocal laser scanning microscopy. The two-color overlay images are also exhibited (Merged). Nuclei were stained with DAPI (blue).

**Figure 4.**
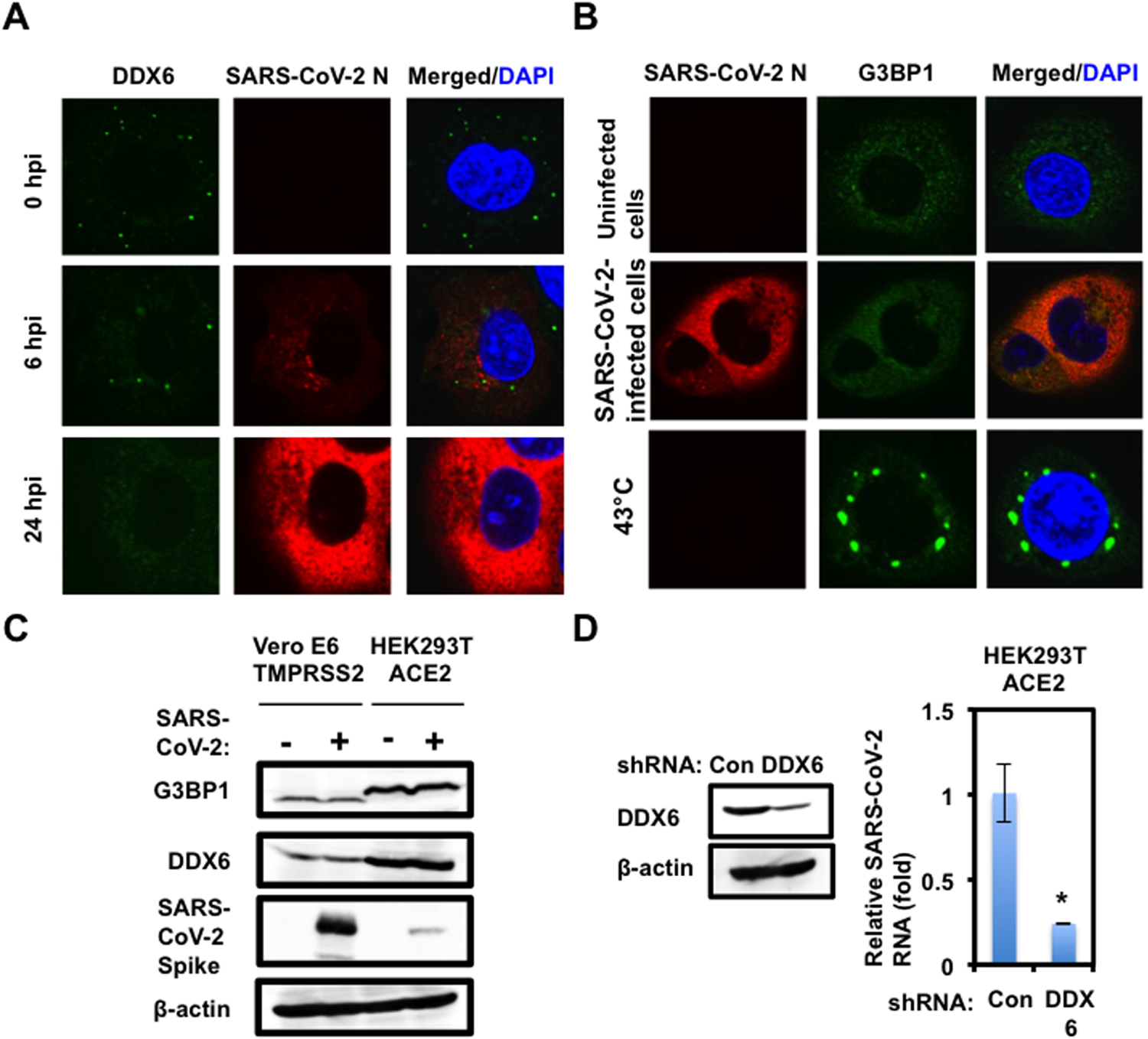
SARS-CoV-2 hijacks DDX6 for own viral replication. (**A**) Dynamic redistribution of DDX6 in response to SARS-CoV-2 infection. VeroE6 TMPRSS2 cells at indicated time (hrs) after inoculation of SARS-C-V-2 were stained with anti-SARS-CoV-2 nucleocapsid (ab273434 [6H3]) and anti-DDX6 (A300-460A) antibodies. (**B**) SARS-CoV-2 does not induce stress granule formation. Uninfected VeroE6 TMPRSS2 or the SARS-CoV-2-infected cells at 24 h post-infection were incubated at 37°C. Uninfected cells were also incubated at 43°C for 45 min. Cells were then stained with anti-SARS-CoV-2 nucleocapsid and anti-G3BP1 (A302-033A) antibodies. (**C**) Host protein expression levels in response to SARS-CoV-2 infection. The results of the Western blot analysis of cellular lysates with anti-SARS-CoV-2 Spike (GTX632604 [1A9]), anti-DDX6, anti-G3BP1, or anti-β-actin antibody in the SARS-CoV-2-infected VeroE6 TMPRSS2 or the HEK293T ACE2 cells at 24 h post-infection at an MOI of 0.5 as well as in the uninfected cells are shown. (**D**) Requirement of DDX6 for SARS-CoV-2 infection. Inhibition of endogenous DDX6 protein expression by the shRNA-producing lentiviral vector. The results of Western blot analysis of cellular lysates with anti-DDX6 or anti-β-actin antibody are shown. The level of intracellular SARS-CoV-2 RNA in the cells at 72 h post-infection at an MOI of 0.5 was monitored by real-time LightCycler RT-PCR. Results from three independent experiments are shown. The level of SARS-CoV-2 RNA in the DDX6 knockdown cells was calculated relative to the level in HEK293T ACE2 cells transduced with a control lentiviral vector (Con). Asterisk indicates significant differences compared to the control cells. *P<0.05.

We next examined whether or not SARS-CoV-2 infection could affect the stress granule formation of G3BP1. The stress granule component G3BP1 dispersed in the cytoplasm at 37°C, while G3BP1 formed stress granules in response to heat shock at 43°C for 45 min in uninfected naïve VeroE6 TMPRSS2 cells (Figure 4B). In contrast, stress granules were not formed in the SARS-CoV-2-infected cells at 24 h post-infection (Figure 4B), suggesting that SARS-CoV-2 infection dose not induce or suppresses stress granule formation. Accordingly, Western blot clearly demonstrated that the SARS-CoV-2 infection did not affect the level of endogenous DDX6 and G3BP1 protein expression (Figure 4C), indicating that SARS-CoV-2 does not induce the degradation of DDX6 and G3BP1. To investigate the potential role of DDX6 in SARS-CoV-2 infection, I examined lentiviral vector-mediated RNA interference to stably knockdown DDX6 in HEK293T ACE2 cells. Western blot analysis for DDX6 demonstrated a very effective knockdown of DDX6 in HEK293T ACE2 cells transduced with lentiviral vectors expressing shRNA targeted for human DDX6 (Figure 4D). Importantly, shRNAs did not affect cell viabilities (data not shown). We next examined the level of intracellular SARS-CoV-2 RNA in the DDX6 knockdown cells 72 h after SARS-CoV-2 infection at an MOI of 0.5. The result showed that the accumulation of SARS-CoV-2 RNA was significantly suppressed in DDX6 knockdown cells (Figure 4D), indicating that DDX6 is required for SARS-CoV-2 infection. Thus, these results suggested that SARS-CoV-2 disrupts the P-body formation of DDX6 and then hijacks DDX6 for own viral replication.

### SARS-CoV-2 N disrupts the P-body formation and hijacks host cellular RNA helicases

Finally, I examined whether or not SARS-CoV-2 nucleocapsid (N) protein interacts with host cellular RNA helicases and affects their subcellular localization, since SARS-CoV-2 N protein is known to bind to the SARS-CoV-2 genomic RNA (32, 33). The SARS-CoV-2 N protein predominantly localized in cytoplasm. I also noticed that SARS-CoV-2 N partially localized in nucleus as well as in nucleoli in 293T cell transfected pcDNA3.1 SARS-CoV-2 N (38) (Figure 5A and 5D). Interestingly, SARS-CoV-2 N disrupted P-body formation of endogenous and ectopically expressed DDX6 (Figure 5A and 5B), suggesting that SARS-CoV-2 N interacts with DDX6. Indeed, both endogenous and ectopically expressed DDX6 dispersed in the cytoplasm and colocalized with SARS-CoV-2 N in 293T cell transfected pcDNA3.1 SARS-CoV-2 N (Figure 5A and 5B). As well, SARS-CoV-2 N disrupted the P-body formation of HA-tagged MOV10 and colocalized in cytoplasm in 293T cells co-transfected with pcDNA3.1 SARS-CoV-2 N and pcDNA3-HA-MOV10 (Figure 5C). Notably, endogenous DDX21 partially colocalized with SARS-CoV-2 N in nucleoli (Figure 5D), indicating that DDX21 interacts with SARS-CoV-2 N. Furthermore, SARS-CoV-2 N colocalized with HA-tagged DDX1 when both proteins were co-expressed in 293T cells (Figure 5E). In contrast, SARS-CoV-2 N did not colocalize with HA-tagged DDX3 (Figure 5F). Therefore, these results suggested that SARS-CoV-2 N interacts with DDX6, DDX21, and MOV10 RNA helicases. To examine whether or not SARS-CoV-2 N binds to DDX21, DDX6, and MOV10, the lysates of either SARS-CoV-2-infected VeroE6 TMPRSS2 cells (Figure 6A) or 293T cells transfected with pcDNA3.1 SARS-CoV-2 N (Figure 6B) were immunoprecipitated with anti-SARS-CoV-2 N, anti-DDX21, anti-DDX6 or anti-MOV10 antibody, followed by Western blotting with anti-SARS-CoV-2 N, anti-DDX21, anti-DDX6 or anti-MOV10 antibody. Consequently. SARS-CoV-2 N were co-immunoprecipitated with endogenous DDX21, DDX6, and MOV10, in both SARS-CoV-2-infected VeroE6 TMPRSS2 cells (Figure 6A) and SARS-CoV-2 N-expressing 293T cells (Figure 6B), indicating that SARS-CoV-2 N binds to DDX21, DDX6, and MOV10. Altogether, SARS-CoV-2 N seems to interact and hijack several host cellular RNA helicases.

**Figure 5.**
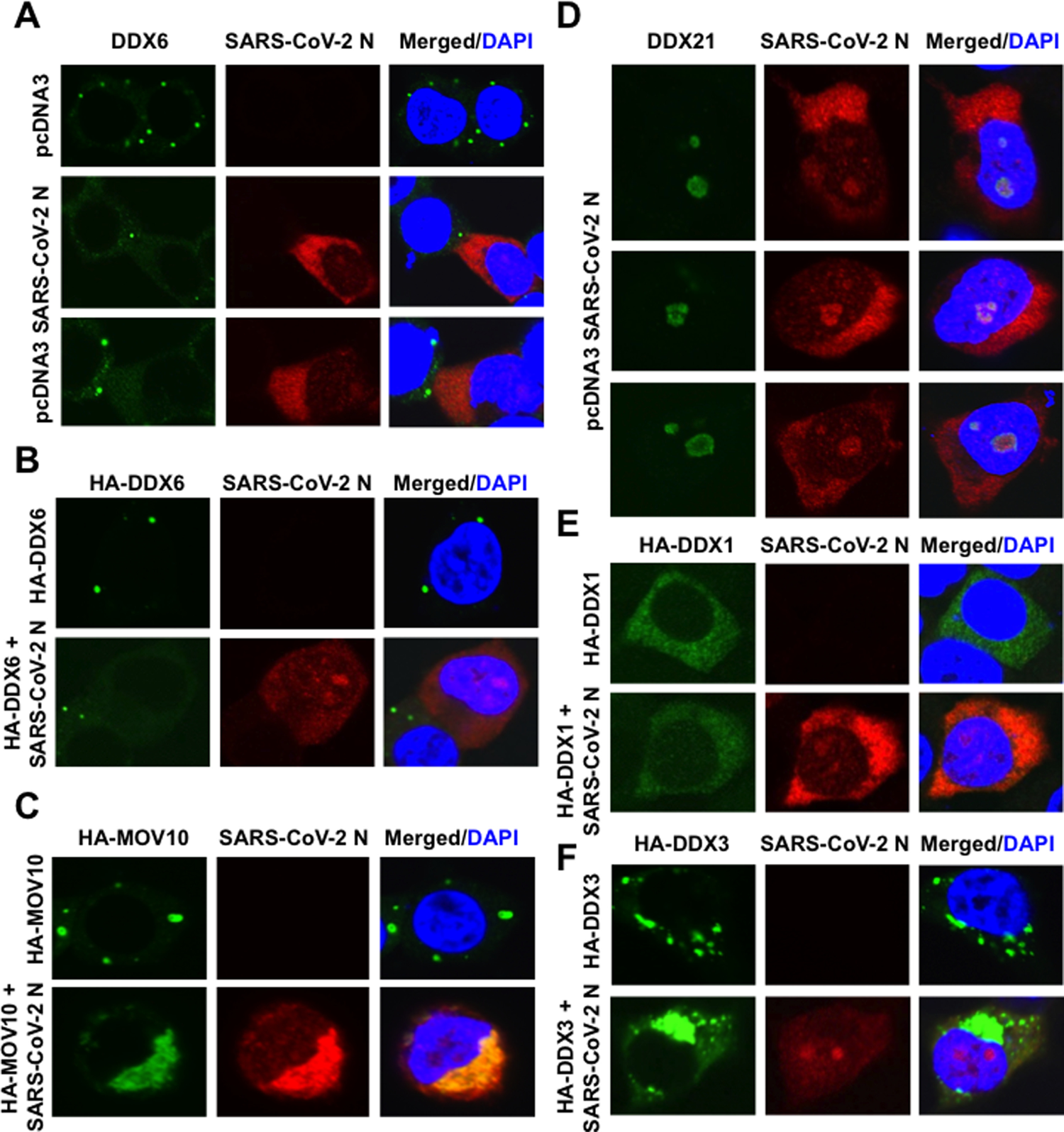
SARS-CoV-2 nucleocapsid (N) protein disrupts the P-body formation of DDX6 and MOV10. (**A**) Disruption of P-body formation of endogenous DDX6 by ectopically expressed SARS-CoV-2 N. 293T cells transfected with 200 ng of pcDNA3.1-SARS-CoV-2 N (38) were stained with anti-SARS-CoV-2 nucleocapsid (GTX632269 [6H3]) and anti-DDX6 (A300-460A) antibodies. (**B**) Disruption of P-body formation of HA-tagged DDX6 by ectopically expressed SARS-CoV-2 N. 293T cells co-transfected with 200 ng of pcDNA3-HA-DDX6 (9) and either 200 ng of pcDNA3.1-SARS-CoV-2 N or pcDNA3 were stained with anti-SARS-CoV-2 nucleocapsid and anti-HA (3F10) antibodies. (**C**) Disruption of P-body formation of HA-tagged MOV10 by SARS-CoV-2 N and colocalization of SARS-CoV-2 N and HA-MOV10. 293T cells co-transfected with 200 ng of pcDNA3-HA-MOV10 and either 200 ng of pcDNA3.1-SARS-CoV-2 N or pcDNA3 were stained with anti-SARS-CoV-2 nucleocapsid and anti-HA antibodies. (**D**) Colocalization of endogenous DDX21 and ectopically expressed SARS-CoV-2 N in nucleoli. 293T cells transfected with 200 ng of pcDNA3.1-SARS-CoV-2 N were stained with anti-SARS-CoV-2 nucleocapsid and anti-DDX21 (A300-627A) antibodies. (**E**) Colocalization of HA-tagged DDX1 and SARS-CoV-2 N. 293T cells co-transfected with 200 ng of pcDNA3-HA-DDX1 (9) and either 200 ng of pcDNA3.1-SARS-CoV-2 N or pcDNA3 were stained with anti-SARS-CoV-2 nucleocapsid and anti-HA antibodies. (**F**) Subcellular localization of HA-tagged DDX3 and SARS-CoV-2 N. 293T cells co-transfected with 200 ng of pHA-DDX3 (8–12) and either 200 ng of pcDNA3.1-SARS-CoV-2 N or pcDNA3 were stained with anti-SARS-CoV-2 nucleocapsid and anti-HA antibodies.

**Figure 6.**
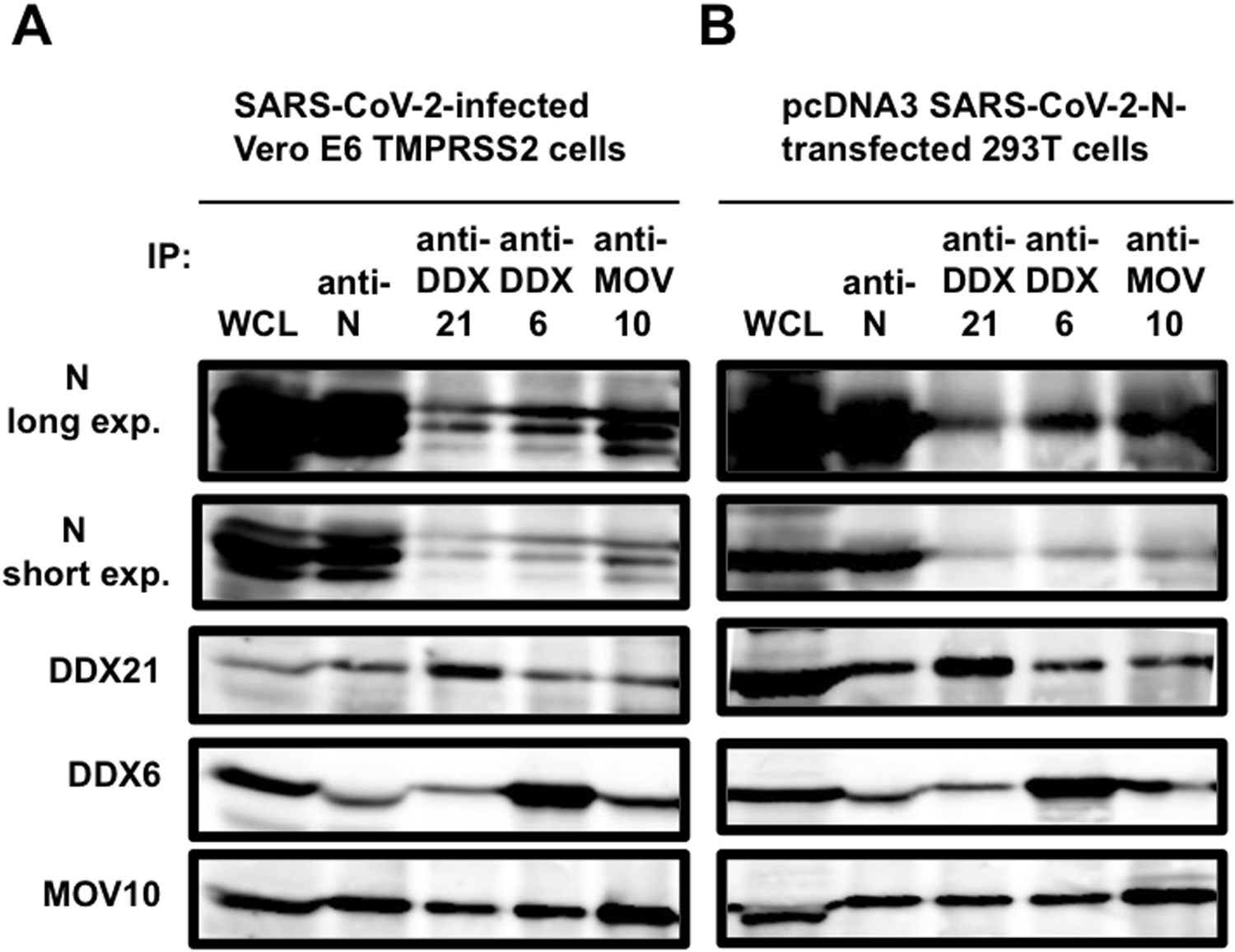
SARS-CoV-2 N binds to DDX21, DDX6, and MOV10. (**A**) VeroE6 TMPRSS2 cells (5×10^5^ cells/well) were infected with SARS-CoV-2 at an MOI of 0.5. The cell lysates were collected at 24 h post-infection. (**B**) 293T cells (2X10^5^ cells/well) were co-transfected with 4 μg of pcDNA3.1-SARS-CoV-2 N. The cell lysates were immunoprecipitated with anti-SARS-CoV-2 nucleocapsid (GTX632269 [6H3]), anti-DDX21 (A300-627A), anti-DDX6 (A300-460A), or anti-MOV10 (A301-571A) antibody, followed by immunoblotting analysis using anti-SARS-CoV-2 nucleocapsid, anti-DDX21, anti-DDX6, and anti-MOV10 antibodies, respectively. Both short exposed and long exposed images were shown.

## Discussion

Several RNA viruses are known to carry their own RNA helicases to facilitate the replication of their viral genome, including alphavirus, coronavirus, flavivirus, HCV, and rubella virus, however, HIV-1 does not carry own RNA helicase (44, 45). Actually, SARS-CoV-2 encodes Nsp13 RNA helicase. Nsp13 RNA helicase is conserved in all coronavirus, including SARS-CoV, SARS-CoV-2, and MERS, and is key player for the viral replication. Interestingly, recent review by Squeglia *et al*. proposed the structural similarity of SARS-CoV-2 Nsp13 RNA helicase with human DDX helicases (46). SARS-CoV-2 has largest RNA genome of ∼30kb among RNA viruses. To regulate and maintain such large viral RNA genome, host RNA helicases may involve in SARS-CoV-2 replication.

In this study, I have demonstrated that several host cellular RNA helicases, including DDX1, DDX5, DDX6, DDX21, and MOV10, facilitate or restrict SARS-CoV-2 infection. Indeed, the DDX21 knockdown markedly accumulated intracellular viral RNA in the SARS-CoV-2-infected HEK293T ACE2 cells (approximately 4,000-fold) compared with that of control cells (Figure 1B), suggesting that DDX21 strongly restricts SARS-CoV-2 infection. The SARS-CoV-2 N protein binds to the viral RNA genome and plays a multifunctional role in the SARS-CoV-2 viral life cycle from regulation of viral replication and transcription and viral genome packaging to modulation of host cell processes. Recent interactome analysis demonstrated that infectious bronchitis virus (IBV) N binds to DDX1, DDX5, DD6, DDX21, and MOV10 (47). The avian IBV also belongs to the *Coronaviridae* family, causes avian infectious bronchitis, a highly contagious disease. IBV N protein localizes in cytoplasm and nucleoli (47). Consistent with this, SARS-CoV-2 N localized in cytoplasm and nucleoli and colocalized with DDX21 in nucleoli (Figure 5D). SARS-CoV-2 N was co-immunoprecipitated with DDX6, DDX21, and MOV10 (Figure 6). Furthermore, recent interactome analysis also supports our finding that SARS-CoV-2 N binds to DDX21 (Figure 6) (48). However, the effect of this interaction on SARS-CoV-2 replication was unknown by only interactome analysis. In this study I clarified the multiple role of DDX21 on the SARS-CoV-2 infection. DDX21 has been involved in multiple functions, including transcription, processing, and modification of pre-rRNA as well as innate immunity. Goodier *et al*. reported that MOV10 associates with the LINE-1 ribonucleoprotein (RNP), along with other RNA helicases including DDX5, DHX9, DDX17, DDX21, and DDX39 (49). Moreover, Zhang *et al*. reported that DDX1, DDX21, and DHX36 helicases form a complex with the adaptor molecule TRIF to sense dsRNA in dendritic cells (50). Thus, the DDX1-DDX21-DHX36 complex participates in innate immunity. Therefore, DDX21 may act as antiviral protein. In fact, it has been reported that DDX21 restricts several viral infection and replication, including influenza virus, dengue virus, borna disease virus (BDV), and human cytomegalovirus (HCMV) (51–54). DDX21 regulates viral replication through various mechanisms, such as suppressing viral genome replication, inhibiting virion assembly and release, and modulating antiviral innate immune responses (51–54). In this regard, I noticed that the DDX21 knockdown accumulated intracellular viral RNA (Figure 1B), the level of extracellular SARS-CoV-2 N protein, and the extracellular SARS-CoV-2 RNA, and elevation of the viral titer and viral infectivity in the supernatants of DDX21 knockdown cells (Figure 2A-2D). Therefore, DDX21 may involve in multiple steps of SARS-CoV-2 life cycle, including intracellular viral RNA replication, viral production, and viral infectivity.

So far, P-bodies and stress granules (SGs) have been implicated in all aspects of RNA life cycle, including RNA translation, RNA silencing, and RNA degradation as well as viral infection (16-18, 26, 27). P body and SG components can facilitate or limit viral infection, and some viral RNAs and viral proteins accumulate into P-bodies and/or SGs. Indeed, P-body components, including DDX6, GW182, Lsm1, and Xrn1 negatively regulate HIV-1 gene expression by preventing viral mRNA association with polysomes (19). In contrast, these miRNA effectors such as DDX6, Lsm1, PatL1, Ago2, positively regulate HCV replication (12, 20–22), since the liver specific and abundant miR-122 interacts with 5’-UTR of HCV RNA genome and facilitates HCV replication (55). In this study, we have found, for the first time, that SARS-CoV-2 infection disrupted the P-body formation of DDX6 and MOV10 RNA helicases (Figure 3). Then, SARS-CoV-2 N colocalized with DDX6 and MOV10 in cytoplasm (Figure 5). Importantly, DDX6 was required for SARS-CoV-2 infection (Figure 4D). In contrast, MOV10 restricted SARS-CoV-2 infection (Figure 1B). On the other hand, SARS-CoV-2 infection did not induce SGs of G3BP1 (Figure 4B). Accordingly, recent interactome analyses indicated that SARS-CoV-2 and IBV N bind to G3BP1 (47, 48). Thus, SARS-CoV-2 infection may attenuate or disturb the SGs formation through the interaction of SARS-CoV-2 with G3BP1. Similarly, we previously demonstrated that HCV hijacks the P-body and stress granule components around lipid droplets (LDs) to utilize own viral replication (7, 12). LDs have been involved in an important cytoplasmic organelle for HCV production (56). In this regard, Dias *et al*. recently reported that SARS-CoV-2 infection up-regulates lipid metabolism and induces LDs, and LDs are required for the production of infectious SARS-CoV-2 viral particles, indicating that LDs are sites for SARS-CoV-2 replication like HCV (57). Therefore, SARS-CoV-2 infection may sequester host cellular RNA helicases around LDs and utilize own viral replication.

Altogether, SARS-CoV-2 seems to hijack host cellular RNA helicases to play a proviral role by facilitating viral infection and replication and, by suppressing host innate immune system. On the other hand, some host cellular RNA helicases including DDX21 and MOV10 limit viral infection and replication as the guardian of host cells.

## Acknowledgements

I thank Drs. Didier Trono, Priscilla Turelli, Hiroyuki Oshiumi, Jeremy Luban, Shutoku Matsuyama, Makoto Takeda, National Institute of Infectious Diseases, and RIKEN BioResource Research Center for reagents. I also thank Dr. Kunitada Shimotohno for valuable discussion. This work was supported by the Research Program on Hepatitis B Grant Number JP17929672 from Japan Agency for Medical Research and Development, AMED.

## Author Contributions Statement

Y.A. contributed to the design of experiments, the conduction of experiments, manuscript writing and editing.

